# Regulating population density through antithetic feedback control of cell growth

**DOI:** 10.1101/2025.04.16.648696

**Authors:** Giovanni Campanile, Vittoria Martinelli, Davide Salzano, Davide Fiore

## Abstract

We present a genetic feedback control strategy enabling engineered microorganisms to self-regulate their population density. Our approach leverages a quorum sensing mechanism for the production of a growth inhibitor protein, whose activation is regulated by an embedded antithetic controller. Through mathematical modeling and steady-state analysis, we provide design guidelines to tune the control parameters. Finally, we validate the control architecture performance and robustness via realistic agent-based simulations in BSim. The proposed control architecture guarantees robust regulation of the cell density while relying on fewer constraints on the biological parameters with respect to other solutions presented in the literature.

## I. Introduction

Engineered microorganisms are widely employed in bio-industrial applications, including biopharmaceutical production and biofuel synthesis [1]–[3]. These microorganisms are endowed with synthetic gene networks that enhance their natural capabilities and enable the production of specific compounds [4], [5]. In doing so, cells redirect metabolic resources from growth and base metabolism to the synthesis of the desired product. A key challenge in these systems is therefore balancing cell growth with production efficiency, thus regulating the cell population density to maintain a desired value becomes necessary to maximize the efficiency of the bioproduction process while minimizing the accumulation of unwanted by-products [6], [7]. This is typically achieved through external control architectures where the growth environment is modified by adding chemicals or nutrients, following the directives decided by a control algorithm hosted on an external computer [8]. However, the manipulation of the environment can face scalability issues, as the cost of modifying the environment grows dramatically as the environment size increases.

A promising alternative is engineering cell populations to self-regulate their growth through embedded feedback controllers, which could enable precise and robust growth regulation without the need of modifying the growth environment. Drawing from natural microbial communities, where cells use intercellular communication to sense and adjust their own population density, various solutions have been proposed in the literature. Most approaches involve coupling a *quorum sensing* mechanism with death to enable self-regulation of population density in single-strain or multiple populations consortia, thereby regulating both relative cell numbers and consortium composition [9]–[15]. In mammalian cells, analogous circuits could be engineered using specific orthogonal communication channels [16]. However, in these works the desired population density is integrated within the design of the synthetic gene network, meaning any change in the working conditions requires re-engineering the system. To address this issue, in [17] the genetic controller is realized by means of a tunable expression system (TES), allowing the desired set-point to be dynamically adjusted online. However, the results presented therein rely on several simplifying assumptions that significantly constrain the possible parameter values.

In this letter, we present an alternative embedded feedback control strategy that enables a cell population to self-regulate its growth rate to reach a desired density. By leveraging a quorum sensing mechanism, cells produce a growth inhibitor protein in response to excessive population density. An antithetic feedback controller [18] modulates the inhibitor production based on the concentration of the quorum sensing molecule, allowing precise control over cell numbers. The population density setpoint is embedded in the gene network design, but can be modified for online tuning using external inputs. Based on analytical results and numerical experiments, we demonstrate that using an antithetic motif instead of the TES employed in [17] alleviates the constraints on the choice of parameters, while enhancing robustness. After describing the proposed control architecture, we derive a mathematical model capturing its dynamics and provide a steady-state analysis of the system allowing us to establish a relationship between some tunable biochemical parameters and the desired density. Finally, we validate the static performance and robustness of the system using BSim, a realistic bacterial populations simulator that enables the simulation of cell growth in a closed environment [19].

## II. Feedback control of the population density

The control strategy we develop here allows a cell population to autonomously regulate its density in a growth environment with *limited resources*, e.g. a vial. This is done through an intercellular communication mechanism, as depicted in Figure 1. Specifically, each cell produces a *quorum sensing* molecule, say *Q*, which diffuses into the environment and acts as a measure of the population density. The measure is then compared to a reference value and, based on the error between the two, a growth inhibitor protein *P* is produced, modulating the growth rate of the cells. The control logic is implemented by means of an antithetic motif involving two biochemical species, namely *Z*_1_ and *Z*_2_, that bind together in a sequestration reaction to form an inactive complex *C*. This motif compares the concentration of the quorum sensing molecule *Q* to the reference value *µ* and adjusts the production rate of *P*. When the concentration of *Q* in the environment increases, the production of *P* increases, leading to a decrease in the population density. Conversely, when the concentration of *Q* decreases, the production of *P* is reduced, allowing the population density to increase again. The antithetic motif has been shown to implement an integral feedback controller that ensures a precise and robust tracking of the reference signal [18]. The desired setpoint for the population density is assumed here to be encoded in the basal production rate of *Z*_1_. However, the design can be modified to allow online tuning of the setpoint by means of external inputs (e.g. light stimuli via optogenetics).

**Fig. 1:**
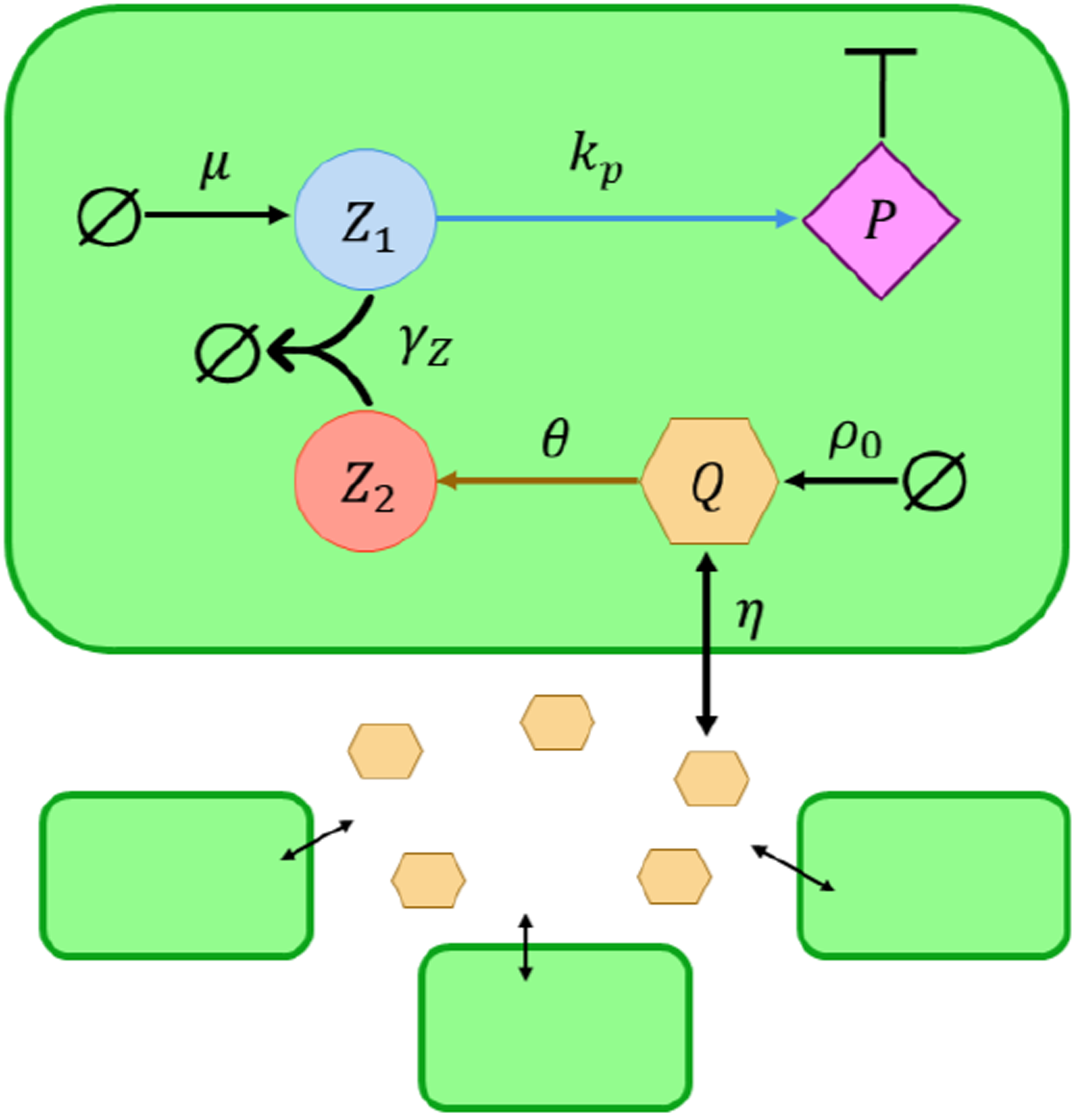
Architecture of the gene regulation network for population density control. *Z*_1_ and *Z*_2_ implement an integral control action through an antithetic motif, by binding together and forming the inactive complex *C. Z*_1_ is constitutively produced at rate *µ*, which encodes the desired set-point 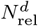, and promotes synthesis of the growth inhibitor protein *P*. *Q* is a *quorum sensing* molecule constitutively produced at rate *ρ*_0_ and broadcasts information about the number of cells in the growth medium by diffusing into the environment and in the other cells; it also modulates production of *Z*_2_, thus closing the feedback loop.

### A. Mathematical modeling

The *aggregate* dynamics of the population can be derived from the single cell model by assuming homogeneity across individuals in the population (see Appendix B). Note that this also implies that all species are diluted at the same rate *γ*.

First, the dynamics of the concentration of the inhibitor protein *P* influencing the population growth rate can be described by:

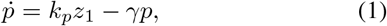

where *k*_*p*_ is the activation rate modeling the strength of the activation induced by the controller species *Z*_1_. The control action is provided by the antithetic motif, where the species *Z*_1_ and *Z*_2_ form an inert complex *C*, and their dynamics is described by:

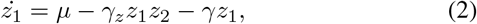

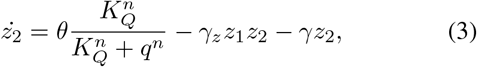

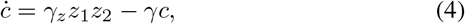

where *z*_1_, *z*_2_ and *c* denote the concentration of the three species. *Z*_1_ is constitutively produced at a rate *µ* which, as it will be shown later, can be tuned to change the desired setpoint. The repression of *Z*_2_ by the quorum sensing molecule *Q* is modeled by a Hill function where *θ* is the production reaction rate, *K*_*Q*_ is the repression coefficient and *n* is the Hill coefficient. Finally, *Z*_1_ and *Z*_2_ bind to form *C* at a rate *γ*_*z*_.

Next, the dynamics of the quorum sensing *Q* within the cells is described by

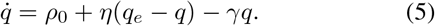

where *ρ*_0_ is its constitutive production rate. In Equation (5), we assumed linear diffusion dynamics across the cell membrane, as often done in literature [20], with *η* being the diffusion rate. Similarly, the evolution of *Q* into the external environment can be written as (see Appendix B for details on the derivation):

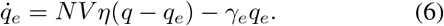

Here, *N* denotes the density of the population growing in the vial, defined as *N* (*t*):= *M* (*t*)*/V*, where *M* (*t*) is the number of cells at time *t* and *V* is the volume of the vial.

Finally, the evolution of the population density is described by a logistic growth model:

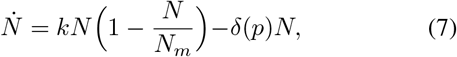

where *k* represents the intrinsic growth rate of the population, and *N*_*m*_ is the carrying capacity, that is, the maximum population density that the limited environment can support. The function *δ*(*p*) represents the growth inhibition rate of the population due to the protein *P*, that we assume to be a linear function of its concentration *p*, i.e. *δ*(*p*) = *d p*, where *d* is the inhibition rate [10].

## III. Control design

In what follows, we establish a relationship between the steady-state value of the population density *N* and the reference parameter *µ*. This relationship provides insights on how variations in the biomolecular parameters affects the steady-state value of population density. To this aim, we first introduce a simplified model of the closed-loop system. Then, via a steady-state analysis, we discuss how changing the biomolecular parameters affects the equilibrium point of the closed-loop system.

### A. Steady-state analysis in fast sequestration regime

To obtain a simple expression of the equilibrium point of the closed-loop system, we start by making the following simplifying assumptions:

**A1** *The Hill function in* (3) *operates in its linear regime*.

**A2** *The sequestration reaction between Z*_1_ *and Z*_2_ *is sufficiently fast*.

**A3** *The quorum sensing molecule diffuses at a rate significantly faster than its dilution*.

#### Remark 1.

*Assumption* ***A1*** *holds if*

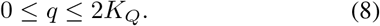

*Under this condition, the repression function of Z*_2_ *due to Q can be replaced with its first-order linear approximation evaluated at q* = *K*_*Q*_, *which simplifies to* 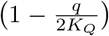 *when the Hill coefficient is n* = 2. *Additionally, Assumption* ***A2*** *implies that the sequestration rate γ*_*z*_ *between Z*_1_ *and Z*_2_ *is sufficiently high, i*.*e, γ*_*z*_ ≫ *γ. Finally, Assumption* ***A3*** *requires that η* ≫ *γ*.

Next, we introduce the change of variables 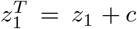 and 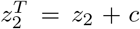, as done in [21], which represent the total concentrations of *Z*_1_ and *Z*_2_ within cells, respectively, accounting for both free and bound forms of these species in the complex *C*. Thus, equations (2)-(4) can be rewritten as

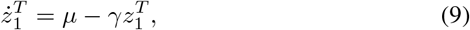

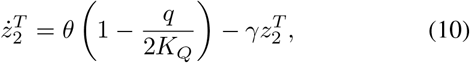

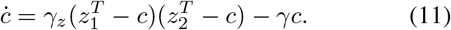

Following [21], we can prove that the trajectories of the system remain bounded within a positively invariant set. To determine this set, we first establish a lower bound. Specifically, when 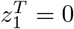, we have 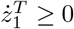. The same holds for all other state variables, ensuring the existence of lower bounds. Next, the upper bound is determined by identifying values of the state variables at which the vector field becomes non-positive. For instance, solving 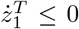 yields 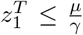. Following the same approach for the other states, we defined the positive invariant set as:

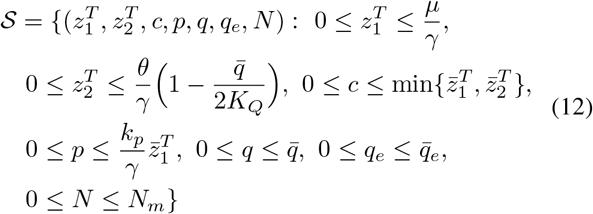

where 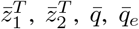 are the steady-state values of 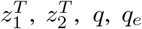, respectively. When the trajectories of the system are bound within the positive invariant set 𝒮 containing the non-trivial equilibrium point, it is possible to derive a relationship between the relative density of the population, that is, the population density *N* normalized by the carrying capacity *N*_*m*_, and the reference input *µ*, as described by the following theorem.

#### Theorem 1.

*Under Assumptions* ***A1****-****A3***, *if the production rate of the quorum sensing molecule Q is set to*

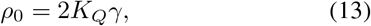

*then, the relationship between the steady-state value of the relative density and the control reference µ can be approximated as:*

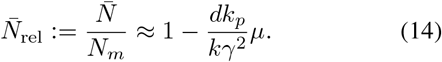

*Proof*. The dynamics of the complex *C* in Equation (11) can be reformulated in a standard singular perturbation form, that is:

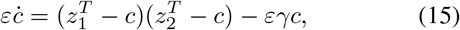

where 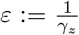 is the perturbing parameter. Under Assumption **A2** we can study the fast sequestration regime by taking *ε →* 0^+^ in (15), in which *c* is the fast variable [21], resulting in

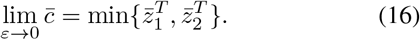

The solution 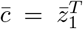 does not correspond to the desired working condition and it will be excluded. Indeed, in this case species *Z*_1_ would be entirely bound in the complex, meaning that there would be none promoting the inhibitor protein *P*, which in turn implies that the control loop was open and the population would grow uncontrolled (as demonstrated by the following steady-state expression for *p* and *N*). Thus, when working correctly, the antithetic motif guarantees that 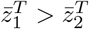, and thus 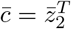.

Setting the time derivatives to zero in equations (9), (10) and (1), (7) yields the following steady-state expressions of the state variables:

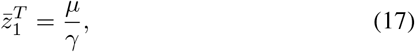

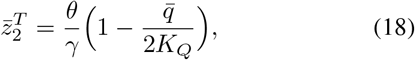

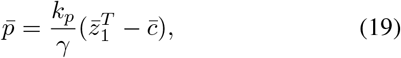

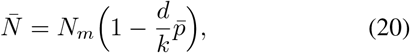

where the steady-state values of *q* and *q*_*e*_ are obtained under Assumption **A3** (see Appendix C for further details) as:

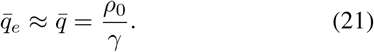

The relationship between the relative density *N*_rel_ and the reference *µ* in (14) can be obtained by first substituting (21) into (18), then (16), (17) and (18) into (19); finally, after the previous substitutions, (19) into (20), yielding the following:

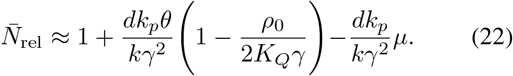

This relationship is a linear function, i.e. 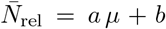 with negative slope *a* = −*dk*_*p*_*/kγ*^2^, and an intercept 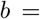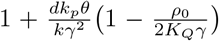. Depending on the value of *b*, three scenarios can occur: (i) if *b <* 1, the controller can regulate the relative density only within subset of [0, 1], namely, [0, *b*]; (ii) if *b >* 1, there is a set of values of 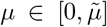 such that 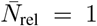, i.e. a saturation effect; (iii) if *b* = 1, the controller can regulate the relative density to any value in the range [0, 1] without saturation, therefore there is a one-to-one correspondence between *µ* and 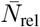. Thus, the range of the achievable steady-state relative density is maximized when *ρ*_0_≤2*K*_*Q*_*γ*. Therefore, by setting *ρ*_0_ = 2*K*_*Q*_*γ* relationship (22) simplifies to (14), thus concluding the proof. ◼

Equation (14) can be recast to derive the expression of the reference parameter *µ* required to achieve the desired set-point 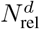:

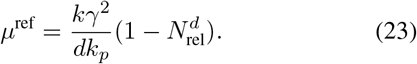

Moreover, equation (14) provides insights on how the biological parameters affect the control system. For instance, if the dilution rate *γ* increases, the term 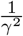 decreases, reducing the sensitivity to the reference parameter *µ*. Additionally, the production rate of the inhibitor protein *k*_*p*_ significantly affects the static gain, and this must be taken into account when choosing the specific protein species in the network.

Note that the structure of (23) is similar to the one presented for the TES in [17] describing the relationship between control input and the desired relative density; however, the implementation presented here employing the antithetic motif requires less constraints on the parameters.

We validated the theoretical findings through numerical simulations in Matlab, see Fig. 2. Here we show that the designed circuit can achieve a wide range of desired steady-state densities.

**Fig. 2:**
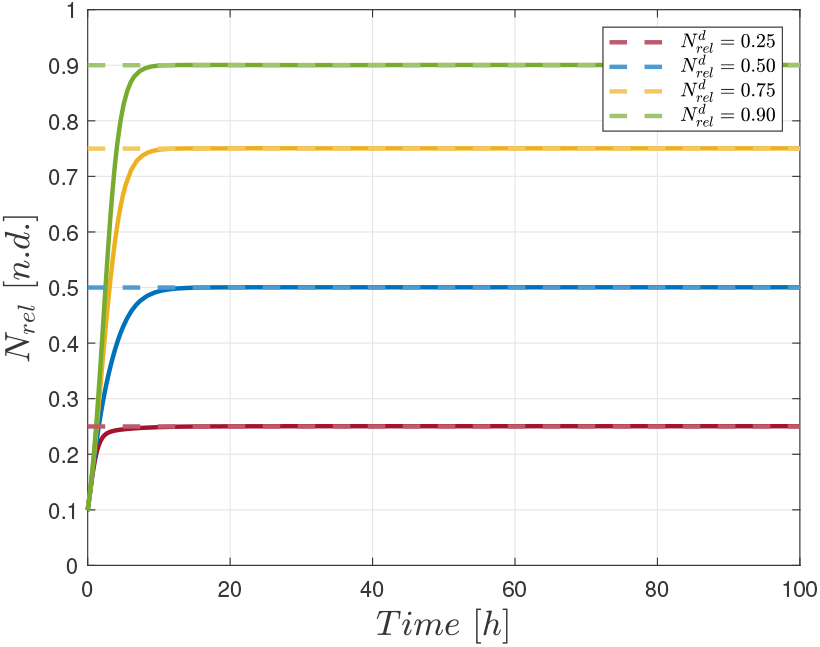
Matlab numerical validation on the aggregate model for several values of the desired relative density. Specifically, 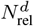 was set to 0.25 (red line), 0.50 (blue line), 0.75 (yellow line) and 0.90 (green line). By tuning the reference parameter *µ* according to Equation (23), the gene network regulates the relative density to the desired set-point 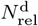. Nominal biochemical parameters are reported in Appendix A.

## IV. In silico experiments

We validated our control strategy in BSim [19], a realistic agent-based simulator for microbial populations that accounts for a variety of effects, including cell-environment interactions, cell growth and replication, and cell-cell communication through chemical species diffusing in the growth medium, e.g. the *quorum sensing* molecules.

In our experiments, we considered a chamber of dimensions 10 *µ*m × 10 *µ*m × 10 *µ*m, closed on all sides, hosting a maximum of approximately 145 cells. These dimensions were chosen as a trade-off between minimizing the computational cost of the simulations and maintaining a sufficient number of cells to preserve statistical significance. Additionally, we modified the numerical routine governing the elongation and division of the cells by modifying the growth rate *k*_*L*_ based on the concentration of protein *P* and the population density *N*, as done in [17].

First, we performed some experiments under nominal conditions, meaning that the biochemical parameters were set to their nominal values (see Appendix A). In this setting, we validated the proposed genetic controller by using several values of the desired relative population density, i.e.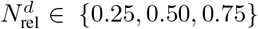. The corresponding values of the reference parameter *µ* was set according to (23). For each value of the desired set-point, the chamber was initially populated with *M* = 10 randomly arranged cells, and the concentration of the chemical species was initialized at zero. Fig. 3 shows both the open-loop (blue line) response and the closed-loop behavior for each set-point, alongside snapshots of the population in the experimental chamber in each scenario. The results confirmed that the proposed controller can regulate the density of the population to the desired set-point, by dynamically adjusting the growth rate of the cells.

**Fig. 3:**
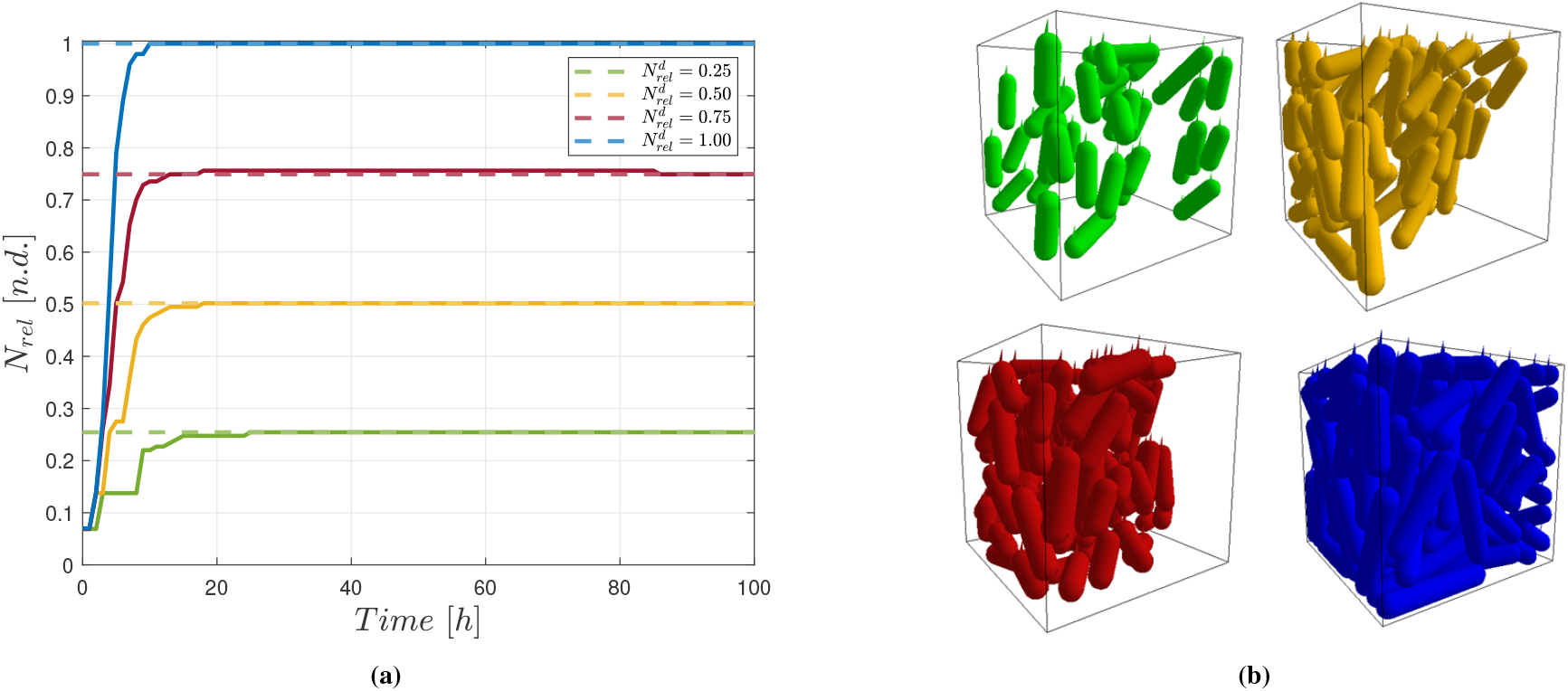
*In silico* agent-based numerical experiments in BSim. (a) Regulation of the relative density *N*_rel_ to several values of the desired set point: 0.25 (green line), 0.50 (yellow line), 0.75 (red line) and open-loop response (blue line). (b) Snapshots of the experimental chamber at steady-state, in which the cells’ color matches the corresponding response in the left panel. Cells are assumed to be identical, and their parameters were set to the nominal values. All simulations were initialized with a population of *M* = 10 cells arranged randomly inside a chamber of dimensions 10 *µ*m × 10 *µ*m × 10 *µ*m. Initial value of the concentration of the chemical species was set to zero. Nominal biochemical parameters are reported in Appendix A.

### A. Robustness to disturbances and cell-cell variability

We tested the robustness of the proposed genetic controller in different settings. First, we considered perturbations consisting in sudden addition and removal of cells in the experimental chamber. Fig 4a shows the relative density when, at times *t* = 50 h and *t* = 100 h, 15 cells, corresponding to approximately 10% of *N*_*m*_, are suddenly added to and removed from the chamber, respectively. By dynamically adjusting the growth rate, cells restore the desired relative density, with a steady-state percentage error equal to 0.38% of the set-point, set to 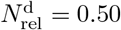.

**Fig. 4:**
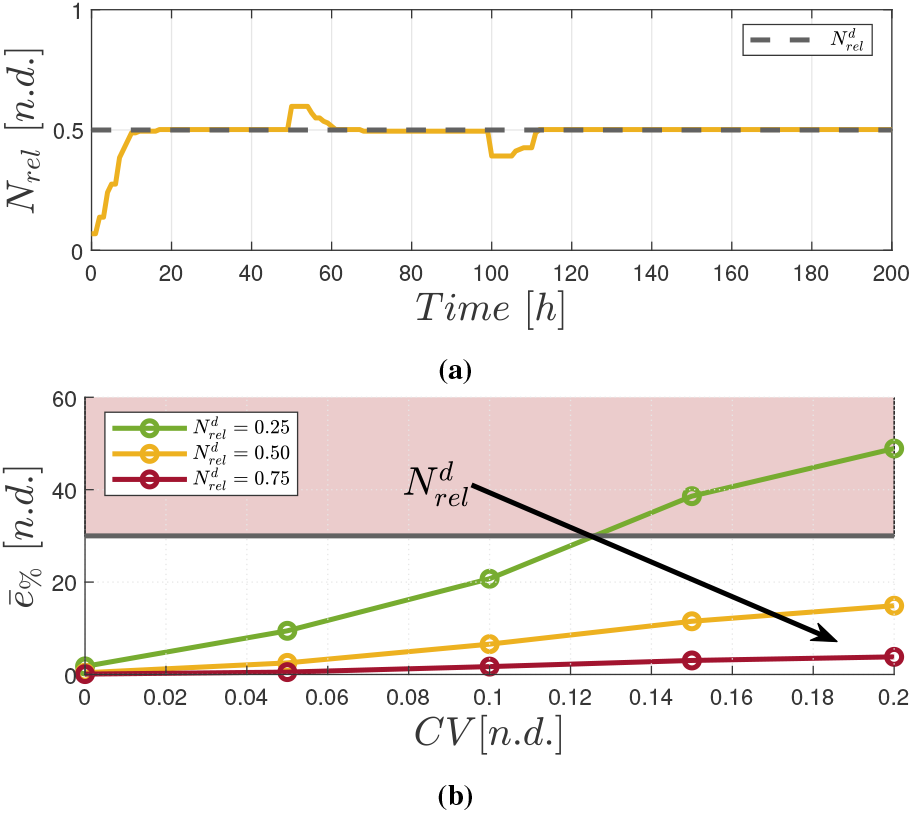
*In silico* robustness experiments in BSim. (a) Robustness to disturbances in the population density. At *t* = 50 h and at *t* = 100 h, 15 cells are added to and removed from the experimental chamber, respectively. The proposed genetic circuit restored the desired relative density 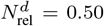 with a steady-state percentage error *e*_%_ = 0.38%. (b) Robustness to intercellular variability. Each parameter, say *α*, was drawn independently from a Gaussian distribution centered in the nominal value, 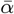, and with standard deviation 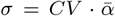. For each couple of values of *CV* and 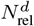 we performed 20 experiments and evaluated the mean steady-state percentage error, considering *e*_%_ *<* 30% to be acceptable in practice. For 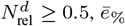 is lower than the threshold for all values of the *CV*. When 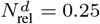, the average steady-state percentage error resulted to be below the threshold for values of the parameters closer to the nominal ones. All experiments were initialized with a population of *M* = 10 cells arranged randomly inside a chamber of dimensions 10 *µ*m × 10 *µ*m × 10 *µ*m. Initial value of the state variables was set to zero.

Then, we tested the robustness to cell-cell variability, modeled in BSim by assigning different parameters to the daughter cells upon division from their mother. Specifically, the parameters we varied in the agent-based model were *θ, k*_*p*_ and the death rate *d*, which affects how the inhibitor protein *p* acts on the population dynamics. Each of these parameters, say *α*, was drawn from a Gaussian distribution centered in the nominal value, 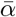, and with standard deviation 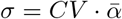, where *CV* is the coefficient of variation.

We considered increasingly larger values of the coefficient of variation, namely *CV* ∈ {0.05, 0.1, 0.15, 0.2}, and several values of the desired set-point, specifically 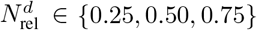. For each pair of values of *CV* and 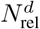, we carried out 20 experiments, and evaluated the average percentage error at steady-state, defined as:

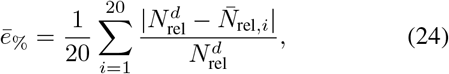

where 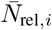 is the relative density at steady-state in the *i*-th experiment.

Fig. 4b shows the results of the robustness experiments, where we considered acceptable a steady-state percentage error below the threshold of 30%. The proposed controller guarantees robustness to cell-to-cell variability in all conditions for values of the desired set-point greater than or equal to 0.50, where the largest average steady-state error is lower than 20%. Below this set-point, the genetic controller proved to be robust when the parameters are closer to their nominal values (low *CV*), while the error exceeded the threshold for larger values of the *CV*. Compared to the TES in [17], the antithetic approach exhibited inferior percentage error at steady-state for all values of the relative density that we considered.

## V. Conclusions

We presented a genetic feedback control strategy that combines a quorum sensing mechanism with an embedded antithetic controller to enable precise regulation of engineered cell population density in a limited culture environment. Our approach, which could potentially improve robustness and scalability of bioproduction systems, offers a promising alternative to other architectures proposed in the literature; for instance, compared to the TES, regulating cell population density with an antithetic controller poses fewer constraints on the system’s parameters, while simultaneously exhibiting better robustness to intercellular variability across a wide range of values for both the relative density and coefficient of variation. However, key challenges still remain, such as defining the correct reference value to achieve the desired set-point. Indeed, while our approach assumes homogeneous cells, natural variability in real populations hinders the selection of a single reference. Additionally, integrating the growth-regulating circuit with other synthetic circuits designed to control the expression of different proteins for various applications could impose an additional metabolic burden on the host cells. Nevertheless, our strategy offers an effective alternative to external feedback control loops and paves the way for communication-based control strategies for maintaining long-term coexistence of different cell populations in microbial consortia [22].

## Appendix

### A. Nominal Biochemical Parameters

The nominal biochemical parameters used in the Matlab and BSim experiments were chosen as follows: *θ* = 18 h^−1^ (taken from [23]); *k*_*p*_ = 5 h^−1^, *k* = 0.97 h^−1^, *η* = 120 h^−1^, *K*_*Q*_ = 20 nM, *n* = 2 and *γ* = *γ*_*e*_ = 2 h^−1^ (taken from [17]); *γ*_*z*_ = 60 (nM · h)^−1^, *d* = 10^−4^ (nM · h)^−1^, *V* = 10 × 10 × 10 *µ*m^3^, *N*_*m*_ = 145*/V µ*m^−3^, and *ρ*_0_ = 2*K*_*Q*_*γ*.

### B. Derivation of the aggregate model

We derived the *aggregate* dynamics from the following equations, which describe the dynamics of the chemical species inside the *i*-th cell of the population:

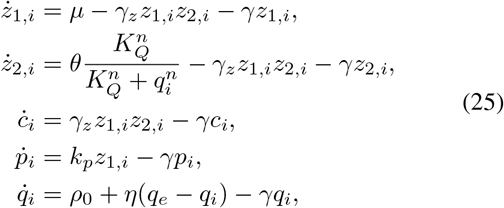

where the parameters have the same meaning as in (1)-(5). By assuming homogeneity between cells in the population, that is, *A*_*i*_ = *A*_*j*_ = *A, A*_*i*_ ∈ {*z*_1,*i*_, *z*_2,*i*_, *q*_*i*_, *p*_*i*_}, ∀*i, j*, where *i* and *j* denote any two cells in the population, equations (1)-(5) immediately follow.

Furthermore, the dynamics of the *quorum sensing* molecule in the external environment, that is, *Q*_*e*_, is described by:

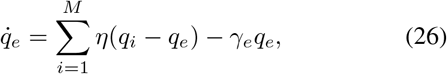

where *M* = *M* (*t*) is the number of cells at time *t*. By expanding the sum in the previous equation, we obtained:

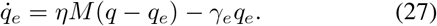

from which equation (6) is derived, by recalling that the population density is defined as *N* (*t*):= *M* (*t*)*/V*, where *V* is the volume of the vial.

### C. Equilibrium point in the fast sequestration regime

To derive the steady-state expression of the *quorum sensing* molecule both inside the cell and in the environment, we first set equation (5) to zero, yielding the following:

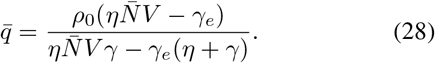

Under Assumption **A3** and considering *γ* = *γ*_*e*_, equation (28) becomes:

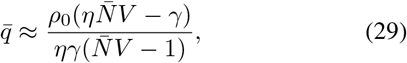

where 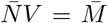 represents the number of cells at steady-state. Since, 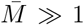 and, as per the previous assumption, *η* ≫ *γ*, it is also 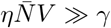. Thus, equation (29) simplifies to 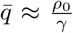. Similarly, by setting the time derivative to zero in (6) and under the same previous assumptions, the steady-state value of *q*_*e*_ is 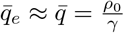.

